# Predicting unknown binding sites for transition metal based compounds in proteins

**DOI:** 10.64898/2026.01.29.702545

**Authors:** Andrea Levy, Ursula Rothlisberger

## Abstract

Transition metal based compounds are promising therapeutic agents, particularly in cancer treatment. However, predicting their binding sites remains a major challenge. In this work, we investigate the applicability of two tools, Metal3D and Metal1D, for this purpose. Although originally trained to predict zinc ion binding sites only, both predictors successfully identify several experimentally observed binding sites for transition metal complexes directly from apo protein structures. At the same time, we highlight current limitations, such as the sensitivity to side-chain conformations, and discuss possible strategies for improvement. This work provides a first step toward establishing a robust computational pipeline in which rapid and low-cost predictors are able to identify putative hotspots for transition metal binding, which can then be refined using more accurate but computationally demanding methods.

**Author summary:** Transition metals play a crucial role as therapeutic agents, especially in cancer therapy. However, the prediction of their binding site locations is challenging, as accurate computational methods often require time-consuming simulations, making them impractical when many possible binding sites must be explored.

In this work, we explored the capability of two binding site predictors, originally developed to locate metal ions in proteins, to identify binding sites for more complex covalently-bound transition metal based agents. We found that these tools can often identify the experimentally-known binding regions, even when starting from the apo structure, in which the protein does not already contain the metal compound. At the same time, our results show clear limitations in more challenging cases, particularly when the binding involves only a single amino acid or when the binding site undergoes major structural rearrangements upon binding.

Overall, our study shows that fast predictors can provide valuable early insights in the investigation of the binding sites of covalently-bound transition metal based compounds. When combined with more accurate simulation techniques, they can help focus computational efforts and ultimately support the rational design of transition metal based drugs.

## 1 Introduction

Compounds based on transition metals (TMs) are promising candidates for biomedical applications, as the combination of organic ligands with TM centers enhances key properties such as stability, solubility, and bioavailability.^1,2^ However, some TMs pose considerable health hazards, and TM-based drugs often show severe side effects. This is the case for cisplatin, which, despite its several toxic side effects, remains a chemotherapeutic workhorse for various tumors.^3^ Significant efforts have been devoted to the search for novel chemotherapeutic strategies to overcome these shortcomings, often relying on alternative TM centers with lower toxicity.^1^

From a theoretical point of view, modeling TM compounds is particularly challenging due to some practical considerations. Drug design often relies on molecular docking,^4,5^ where different binding poses are evaluated with some loss function to identify candidates showing high affinity. However, TM-based compounds can bind biomolecules through the formation of one or more covalent bonds, and in docking, accurately predicting the binding pose of covalent compounds is particularly challenging.^6,7^ In the investigation of drug candidates interacting with specific protein targets, often molecular dynamics (MD) simulations based on classical force fields (FFs) are performed to evaluate the energetics and dynamics of the compound at different putative binding pockets. However, in the case of TMs, classical FFs often lack accuracy and are not always able to reproduce, e.g., the correct coordination geometry.^8^ In addition, standard FFs are not capable of describing the chemical reactions that lead to the formation of covalently or coordinatively bound ligands.

One possible solution to overcome these limitations is to use hybrid quantum mechanics/molecular mechanics (QM/MM) methods, which are able to accurately describe bond formation events by taking the electronic structure of the TM-based complex and its protein environment explicitly into account.^9–11^ However, such methods are orders of magnitude more computationally expensive than molecular docking or even classical FFs approaches, making them impractical to use for high-throughput studies in the case where the binding site for a TM-based compound in a protein is unknown and various putative sites have to be assessed. For this reason, fast but accurate predictors to highlight potential ‘hotspots’ for the binding of TM complexes are fundamental to focus the computational effort where it is really needed.

We recently introduced two prediction tools to locate the binding site of zinc ions in proteins: Metal3D, based on a 3D convolutional neural network (3DCNN), and Metal1D, based on geometric criteria and a probability map associated with the metal under study.^12^ Even if solely trained on zinc, Metal3D is able to accurately locate different TM ions. This is particularly interesting for TM binding site location, since the training of 3DCNNs and machine learning (ML) models in general is computationally expensive and often requires a large amount of data. Metal1D showed similar precision in predictions for zinc and other TM ions, i.e., most predictions made are correct, but lower recall, i.e., a larger number of binding sites are not predicted in the case of TM ions different from zinc. However, since Metal1D relies on probability maps and geometric rules only, it can be expected to be less data-demanding than Metal3D, as well as more easily adaptable to different metals with low computational cost for generating a new probability map.

The ability of Metal3D and Metal1D to predict the binding site of other TM ions triggered the question of whether these tools are also capable of providing useful predictions for the location of binding sites for TM-based compounds. Such compounds present different organic ligands that coordinate the metal center, introducing steric constraints on the binding site accessibility. In addition, the presence of ligands coordinating the metal center limits the number of possible bonds that it can form with protein ligands, typically resulting in one or two coordinating amino acids only. However, binding sites for such compounds and for bare TM ions share several similarities, particularly in the types of amino acid residues involved.

The focus of this paper is to probe the performance of Metal3D and Metal1D for the identification of binding sites for TM-based agents. In the case of Metal1D, we also evaluated binding site predictions performed with two additional probability maps generated for this study, i.e., for Pt- and Ru-based compounds. In the following, Sec. 2 briefly illustrates the details of the two predictors and the systems under study. Results for different protein systems are presented in Sec. 3, while Sec. 4 presents the main conclusions and outlooks of this study.

## 2 Methods

### 2.1 Metal3D and Metal1D predictors

A detailed description of Metal3D and Metal1D can be found in the original publication by Dürr et al. ^12^ Here, we provide a short overview of the main features of the two predictors and highlight their differences and similarities. A schematic representation of their training and inference mode to predict metal ion positions is provided in Fig. 1.

**Figure 1:**
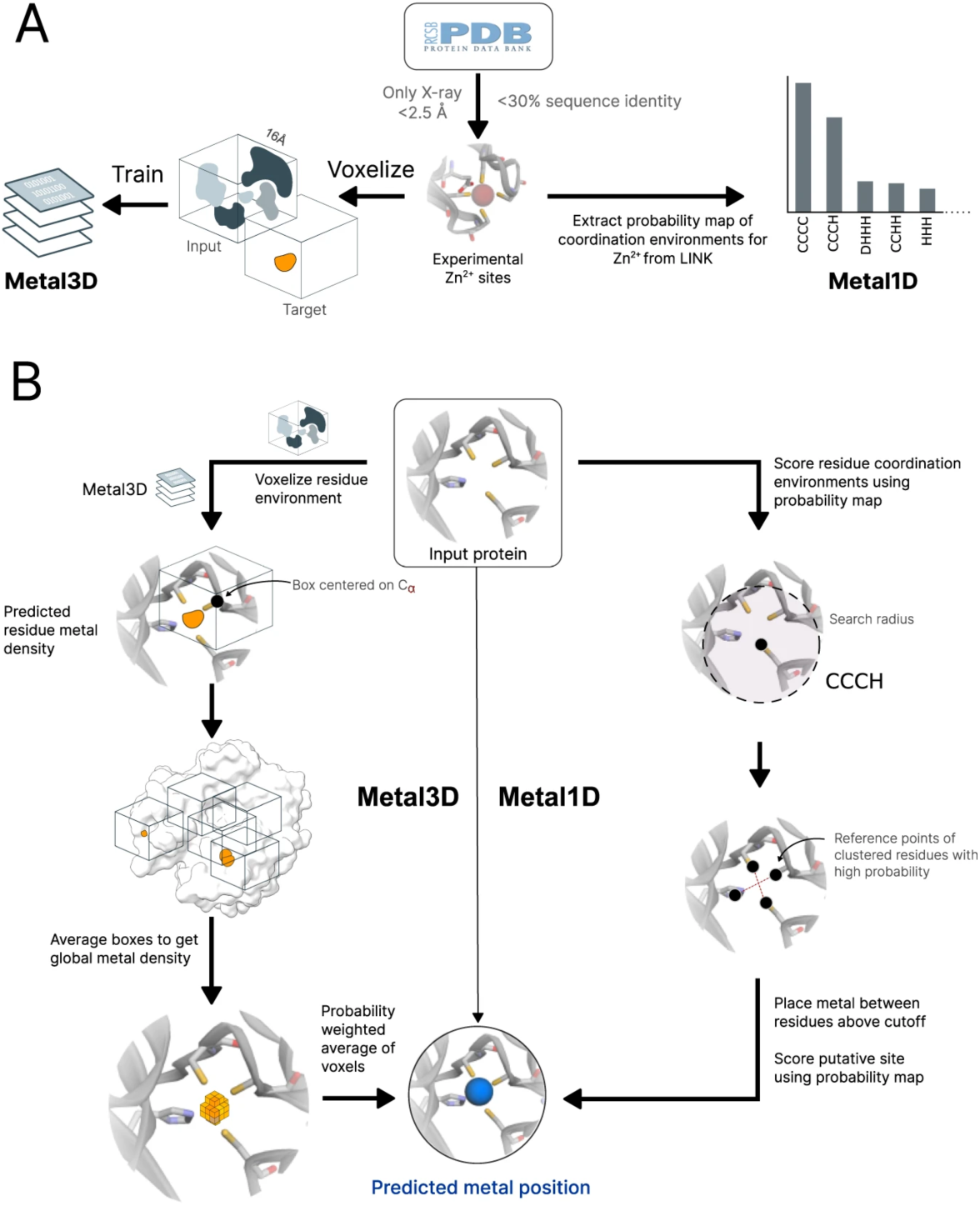
Schematic representation of the training (A) and the inference mode (B) for Metal3D and Metal1D predictors. Reproduced from Ref. 12 under CC BY 4.0 license.

Metal3D is based on voxelization of input structures into three-dimensional grids, which are then fed to a 3DCNN that uses eight different input channels to describe the molecular environment in a voxel, i.e., aromaticity, hydrophobicity, positive/negative charge, hydrogen bond donor/acceptor, occupancy, and metal ion binding side chain. The model uses boxes of size 16 Å centered on the C*α* atom of different residues, with a grid resolution of 0.5 Å, to generate a probability density for metal ion location at each grid point starting from the coordinates of the atoms in the input box. This is done through a series of convolutional layers with filter size 1.5 Å, except for the fifth layer where a larger filter (8 Å) is used to capture interactions on a longer range. The probability density from boxes within 0.25 Å of each grid point is then combined by averaging the probability box densities to generate a global probability for the entire protein, which can be used to place metal ions. However, for this project, we focused on the total probability density itself rather than on the exact placement of the metal. This allowed us to systematically explore a broader range of probability values, including low-probability sites that might otherwise be overlooked when Metal3D does not place a metal ion.

On the other hand, predictions performed with Metal1D are based on a probability map derived from the LINK records in the protein structures from the RCSB Protein Data Bank (PDB),^13^ which specify the connectivity between a ligand (the metal ion in this case) and the amino acids of a protein. To perform a prediction, a geometrical search is performed around each amino acid within a search radius, starting from a reference point corresponding to the atom most likely to coordinate a metal. Based on the surrounding amino acids within the search radius, a score is assigned to the reference point by summing the probabilities in the probability map for coordinations compatible with the one observed. After each amino acid in the protein is scored with this approach, the putative metal sites are predicted by grouping the highest-scored amino acids in clusters based on distance and locating a putative site as the weighted average between the coordinates of the reference point of each amino acid, using the amino acid score as a weighting factor. Each site is then re-scored by performing a geometrical search centered on the site position and assigning a probability score based on the probability map for the given metal, in the same way as the amino acid scores are computed.

### 2.2 Metal binding site predictions

In this study, we chose two systems to test the prediction ability of Metal3D and Metal1D for TM-complexes: hen egg-white lysozyme C (HEWLC, UniProt ID P00698) and bovine pancreatic ribonuclease A (RNaseA, UniProt ID P61823), for both of which experimentally determined binding sites for Pt- and Ru-based compounds are known. For each protein, we selected different crystal structures to create a representative set that includes both the apo protein and holo structures containing Pt- and Ru-based agents. The selection of multiple structures for the same protein serves a dual purpose. First, it enables the identification of experimentally known binding sites to use as a reference. Additionally, it allowed us to evaluate the stability of the predictors with respect to small changes in the input structure: albeit referring to the same protein, different experimental structures can show some variations, e.g., in the side chain positioning or rotameric state, and ideally, a good predictor should have limited sensitivity to such changes. In particular, for the HEWLC enzyme, we selected PDB IDs 194L,^14,15^ 2I6Z,^16,17^ 5II3,^18,19^ 6QEA,^20,21^ 5V4G,^22,23^ and 6WGO,^24,25^ while for the RNaseA enzyme we selected PDB IDs 1RPH,^26,27^ 4S0Q,^28,29^ 4S18,^29,30^ and 5JLG.^31,32^ More details about the similarity between the different protein structures are reported in the corresponding sections, as well as in the Supporting Information (SI).

To test the sensitivity with respect to different rotamers in the case of a bidentate TM-based compound, i.e., binding to two amino acids in the protein, we introduced a third test system extracted from the crystal structure of a nucleosome core particle (NCP) in which the TM complex binding induces a conformational change. We investigated the binding site composed of Glu61 and Glu64 in the H2A histone protein, named GLU site.^33^ In particular, we considered the GLU site from the apo structure (PDB ID 3REH^34,35^) and a second structure in which it is occupied by a bidentate Ru-based compound (PDB ID 5XF3,^19,36^ more details in the corresponding section and in the SI). Upon binding of the metal compound, the side chain conformations of the Glu61 and Glu64 are different. This feature is consistently observed in several crystal structures, e.g., the conformation of the apo structure is the same as in other crystal structures in which the site is not occupied, e.g., in PBD IDs 4J8U^37,38^ and 4J8W,^38,39^ and analogously, the observed rotamer change in the holo structure is shared by PDB ID 5DNN,^40,41^ where a different Ru-based compound is co-crystallized (more details in Fig. S6).

In the case of predictions performed with Metal3D, we analyzed the global probability density file generated using VMD,^42^ which has also been used to compare the RMSD of the different structures. Since we are interested in exploring the possibility of using it for predicting TM-based compounds, and we expect an associated probability significantly lower than the one for a typical zinc binding site, we analysed regions with relatively low probability values by finding the minimal isovalue that enabled localizing distinct sites around the protein (p∼0.04).

In the case of Metal1D, there exists the possibility of selecting a threshold value to exclude predictions that have too low probability scores. To maximize the number of predictions and avoid overlooking any possible site, even with low probability, we set a threshold of 0.99, i.e., all predictions with a probability within 99 % of the highest-scored one are considered. Metal1D then provides the coordinates of putative binding sites and a probability score associated with them. We investigated the site location by visual inspection using VMD.^42^ In addition to predictions performed with the original probability map for zinc ions,^12^ we also introduced two additional probability maps for Pt- and Ru-based compounds. A detailed explanation of which PDB structures have been used to generate these maps is provided in Sec. 3.1. The new probability maps, as well as a Jupyter notebook to generate them, have been deposited on Zenodo under https://zenodo.org/records/18416988.

It is important to note that both tools base their predictions only on the coordinates of the heavy atoms in the protein crystal structure. The hydrogenation or protonation state of the different amino acids is not taken into account explicitly, so the absence or presence of hydrogen atoms does not affect the prediction. Moreover, even if the crystal structure includes non-protein atoms, such as ions, water molecules, or other ligands, they are not considered when performing a prediction.

### 2.3 Rotamer sampling

We explicitly investigated the effect of different rotameric states on the predictions by Metal1D and Metal3D. To generate such conformations, we used an in-house Jupyter Note-book available on Zenodo (https://zenodo.org/records/18416988). To sample different rotamer conformations, we used a similar approach as in the EVOLVE package:^43^ rotamers are derived from the Richardson rotamer library,^44^ which corresponds to the most commonly observed side chain conformations for the naturally occurring amino acids in the PDB, i.e., low-energy conformers of the side chain torsion angles {*χ_i_*}. For each sampled rotamer, the backbone atoms are aligned to those in the original structure, and we performed a separate Metal3D or Metal1D prediction on the resulting conformation.

## 3 Results and Discussion

### 3.1 Differences in data availability

In the case of Metal1D, we generated two additional probability maps to compare them with the results from the original one based on zinc, using PDB structures containing Pt or Ru ions, as well as complexes containing such TMs. The generation of the probability map is analogous to the one for zinc described in Ref. 12, and is based on the LINK records in the PDB structures. In the case of Pt, fewer than 250 crystal structures in the PDB database contain a Pt-based compound covalently bound to a biomolecule. Among those structures, we identified only 81 X-ray structures with resolution higher than 2.5 Å corresponding to unique proteins, containing 244 binding sites, i.e., where Pt ions (or Pt atoms in Pt-based complexes) contain a LINK to at least one amino acid. In the case of Ru, these numbers are even more reduced: about 160 crystal structures with covalently bound Ru-based complexes are present in the PDB, and only 28 unique protein structures, corresponding to 47 Ru binding sites. The number of available structures is drastically low in comparison with what was used to train the 3DCNN model for Metal3D, where more than 2000 crystal structures corresponding to unique proteins with high resolution have been used, featuring ∼15 600 zinc binding sites. This comparison highlights how challenging the problem of designing a predictor for some TM binding sites can be due to the limited amount of data for training, especially in the case of 3DCNNs, and ML models in general.

To compare the binding propensity to different amino acids, we report in Fig. 2 the frequency of binding for the different amino acids extracted from the LINK analysis for the Zn(II) ion used in the original training of Metal1D, as well as the ones resulting from generating the probability maps for Pt- and Ru-based compounds (including ions and metal-based compounds). In the case of Zn(II), the preferential binding amino acids are histidines, followed by cysteines and, at lower frequencies, aspartates and glutamates. This is in line with previous studies in which the PDB has been analyzed to determine the TM binding selectivity in proteins.^45–48^ From this simple comparison, it is possible to observe metal-specific characteristics in the amino acid binding patterns, which will be reflected in the corresponding probability maps for Metal1D. This is particularly evident in the case of Pt-based compounds, where methionine binding becomes preferential and presents even higher probability than other typical metal-binding amino acids.

**Figure 2:**
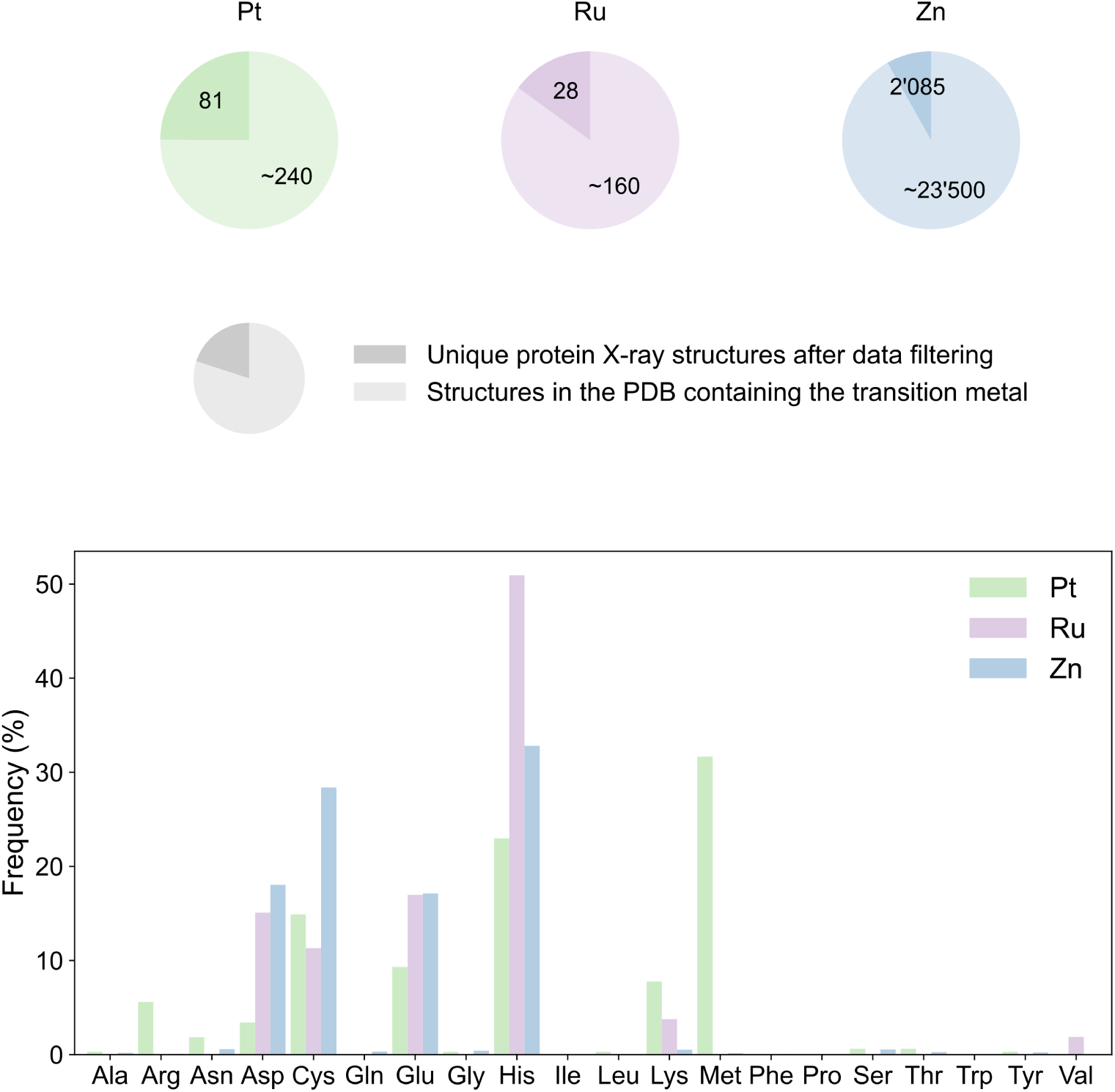
Dataset statistics on the PDB structures used to generate Metal1D probability maps for Zn(II) ions and for Pt and Ru (ions and metallocompounds). The pie charts at the top show the difference in the available number of experimental structures from the PDB for the different cases, and their further reduction after filtering to generate the probability maps for Metal1D. The histograms at the bottom show the binding preference to different amino acids, based on the analysis of the LINK records. Note that for each case, the histogram is generated from the unique protein X-ray structures after data filtering.

### 3.2 Lysozyme C TM binding site prediction

The hen egg-white lysozyme C (HEWLC) protein was the first enzyme for which a three-dimensional structure has been resolved^49,50^ and, to date, is among the enzymes with more crystal structures available thanks to its propensity to easily crystallize in various crystal forms, yielding high-resolution X-ray data.^51^ Our test set (Fig. 3) contains an apo HEWLC structure,^14^ one in complex with cisplatin,^16^ two with different Pt(II)-based compounds binding the protein in a planar geometry,^18,20^ and two different Ru(II)-based compounds at different binding sites.^22,24^ All the TM compounds considered form monodentate adducts, binding a single amino acid.

**Figure 3:**
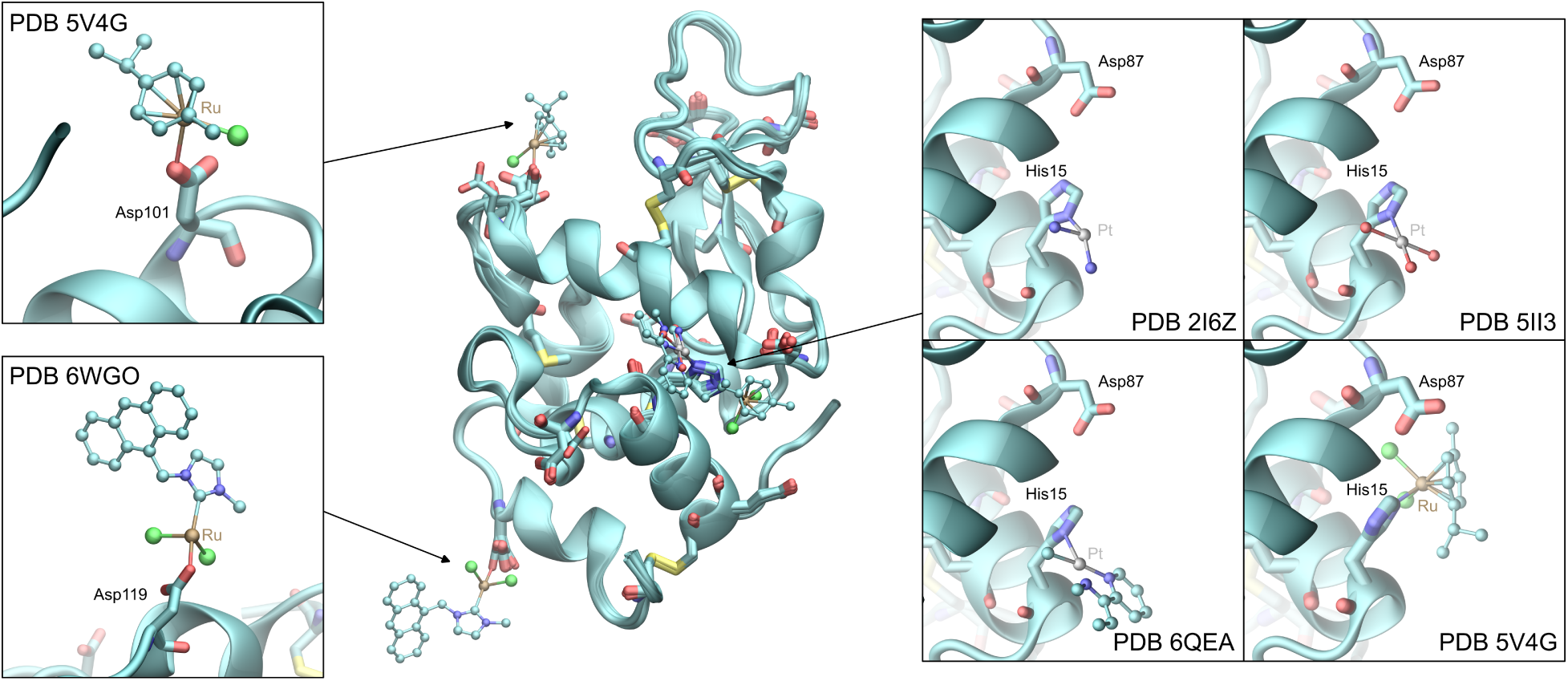
PDB structures for the HEWLC enzyme used as test set. All the PDB structures have been aligned, and the metal binding amino acids are represented in licorice representation. The binding sites for each TM compound are reported in different inserts.

All the holo structures containing TM complexes deviate only slightly from the apo structure with RMSD lower than ∼0.3 Å for the backbone atoms, and ∼0.9 Å when considering the metal binding amino acids (Asp, Cys, Gly, His, and Met), including their side chains. In particular, focusing on the amino acids of the three binding sites considered, the binding of a TM-based compound at His15 is associated with a different rotamer, while in the case of Asp119, only a different positioning of the side chain is observed, without a change of the rotameric state. The binding site at Asp101 appears to be more mobile, with different side chain positioning and rotamer changes observed among the different structures, even in the absence of a binding TM compound. More details about the rotameric states, as well as the RMSD values, are reported in the SI (Table S1).

Starting with the discussion of the results from Metal3D, we notice that the predictions for the different structures are quite consistent, with small changes in the regions with larger probability. The predictions for the apo structure are reported in Fig. 4, while the predictions for all the other structures are reported in Fig. S1. In particular, the region around His15, i.e., the binding site for the Pt-based compounds and one of the Ru-based ones, corresponds to the highest probability region predicted by Metal3D, with values ≥0.10–0.15 depending on the structure. This result is not affected by the change in conformation observed for some of the structures, which corresponds to a rotation of the imidazole ring. This might also be due to the proximity of Asp87, which makes the region between the two amino acids favorable for the binding of TMs. It is important to notice that the probability values are significantly lower than what Metal3D predicts for, e.g., Zn(II) binding sites. This can be attributed to different factors: zinc ions often bind with tetracoordinated geometries with at least four binding amino acids, resulting in higher probability values compared to the Pt and Ru complexes forming mono- or bi-dentate adducts with proteins. Nevertheless, even if with low probability in absolute terms, Metal3D is able to identify the binding site at His15.

**Figure 4:**
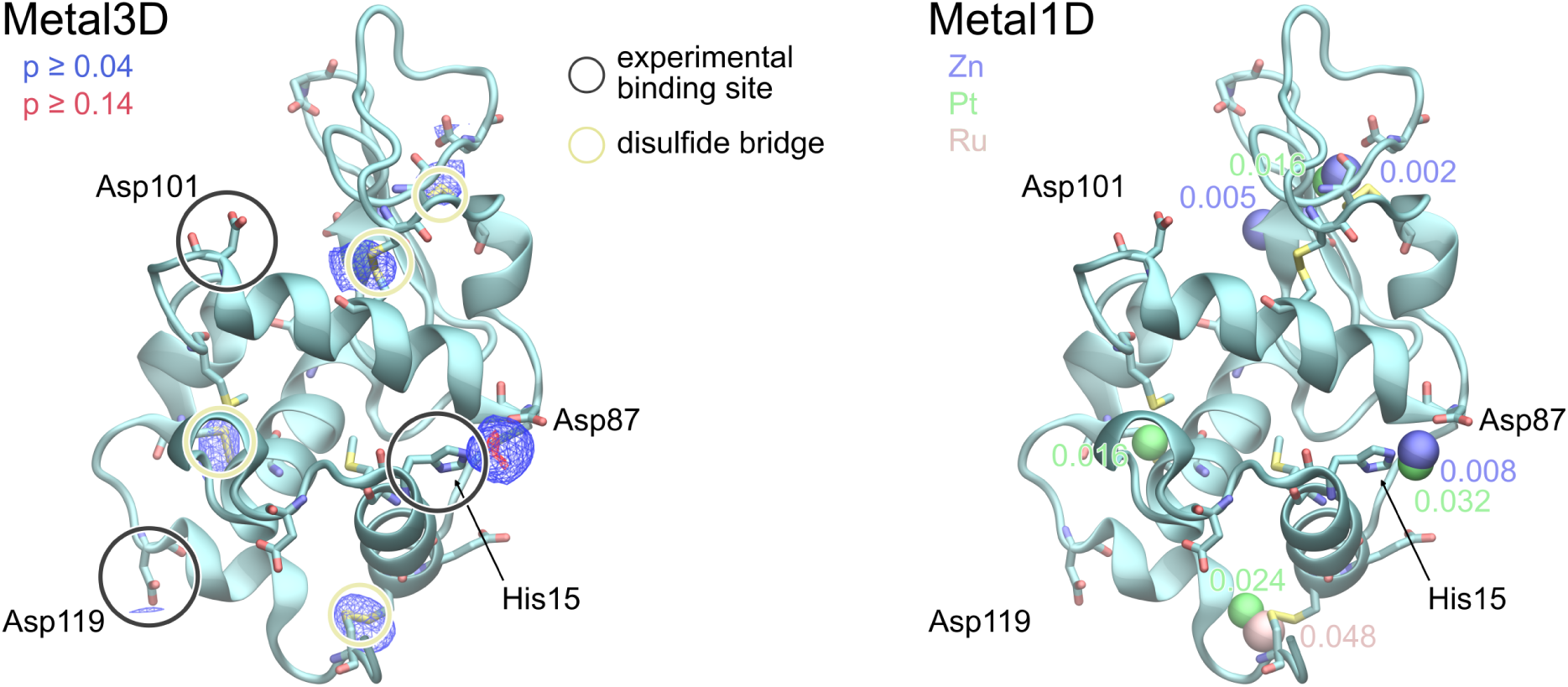
Metal3D (left) and Metal1D (right) predictions for the apo structure of HEWLC (PDB 194L). The experimental binding sites and disulfide bridges are highlighted with circles in the structure on the left. In the test set considered, Ru-based compounds bind all three sites, while only Pt-based compounds bind at His15. For Metal3D, two probability density contours are represented in different colors as isosurfaces with a wireframe representation: lower probability isovalues of 0.04 in blue and higher ones (0.14) in red. For Metal1D, the results obtained with different probability maps are reported: zinc (blue), platinum (green), and ruthenium (pink), and for each prediction, the probability associated with each site is reported in the figure.

Concerning the two other binding sites for Ru-based compounds, both corresponding to a single aspartate residue (Asp101 or Asp119), they are not successfully predicted by Metal3D: some very low-probability region is identified in some of the cases (p∼0.04), but this prediction is not consistently found in all the structures. This is true even in the case of the holo structures, for which only extremely low-probability regions, or no regions at all, are identified at Asp101 or Asp119. This is interesting since it can be assumed that the amino acid side chain in structures containing a TM complex is in ‘optimal’ conformation for binding. However, both sites present a single metal binding amino acid, which is a really different environment from a typical Zn(II) ion binding site, on which Metal3D was trained. Other Metal3D predictions, consistent among all the structures, correspond to the four disulfide bridges present in the protein structure. Considering that cysteines are the second most likely amino acids to bind Zn(II) (Fig. 2), it is not surprising that such regions are identified as probable for metal binding. However, in practice, from visual inspection of the protein structure, it is straightforward to discard such regions when investigating binding sites for TMs.

Also in the case of Metal1D (predictions for the apo structure in Fig. 4 and for all other structures in Fig. S2), the His15 site is identified consistently in all the structures with the highest probability values using the original probability map from Zn(II) structures. This is also the case when using the probability map generated for Pt-based compounds, which, in general, identifies similar sites as the zinc one, including some of the cysteines forming disulfide bridges, similarly to what has been observed in the Metal3D predictions. Moreover, predictions performed with the Pt-based probability map present larger absolute values than the ones from the Zn-based probability map, e.g., with a probability associated with the site at His15 of 0.032 for the Pt map, and 0.008 for the Zn map (in both cases, this is the highest probability site predicted). These values are significantly lower than predictions of Zn(II) tetracoordinated binding sites, but the same considerations as the ones for Metal3D predictions apply. No Metal1D prediction is located in proximity of the two binding sites at Asp101 and Asp119, suggesting that Metal1D, as Metal3D, fails to predict such sites containing a single amino acid ligand. Concerning the prediction performed with the probability map extracted for Ru-based compounds, in all structures, only one or two disulfide bridges are identified, and even if histidines are the most frequent Ru-binding amino acids in the probability map for ruthenium (Fig. 2), no prediction occurs for the His15 binding site.

### 3.3 Ribonuclease A TM binding site predictions

Bovine pancreatic ribonuclease A (RNaseA) is another extensively studied protein, which was the first enzyme to have its complete amino acid sequence determined. ^52,53^ Our test set (Fig. 5) contains an apo RNaseA structure,^26^ two with different Pt(II)-based compounds forming mono- and bi-dentate adducts at different binding sites,^28,30^ and one with a Ru(II)-based monodentate compound.^31^ This second set of protein structures allows us to extend the test of Metal3D and Metal1D in the prediction of binding sites for a larger variety of Pt and Ru compounds, also including a bidentate site. From a practical point of view, such sites are closer to ideal binding sites of TM ions, potentially increasing the possibility of predictions for Metal3D and Metal1D.

**Figure 5:**
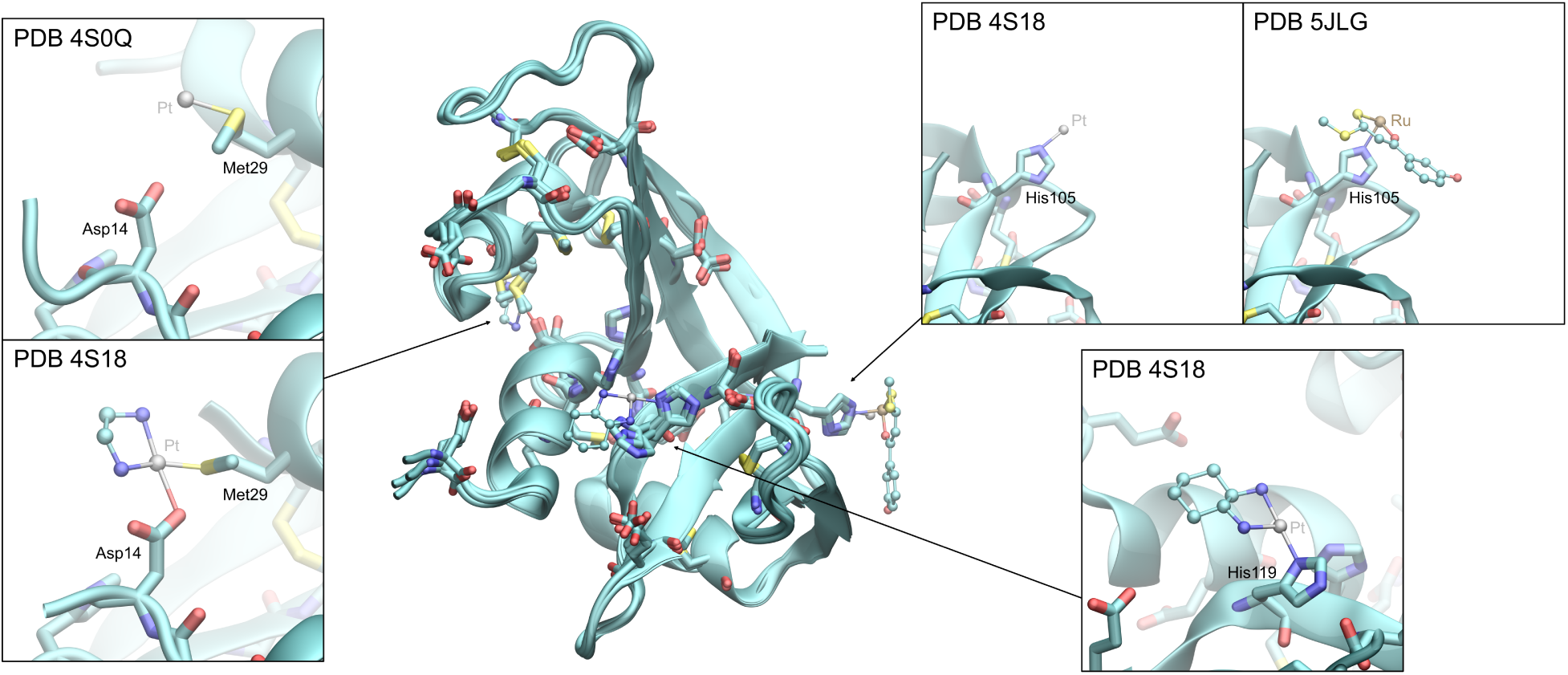
PDB structures for RNaseA used as test set. All the PDB structures have been aligned, and the metal binding amino acids are represented in licorice representation. The binding sites for each TM compound are reported in different inserts.

Similar to HEWLC, the holo structures in the test set are similar to the apo one, enabling us to probe the sensitivity of the predictions to small changes in the side chain conformation. In particular, all the structures deviate from the apo structure by less than ∼0.5 Å RMSD for the backbone atoms, and ∼0.7 Å for the binding amino acids (Asp, Cys, Glu, His, and Met), including their side chains. The binding sites at His105 and His119 do not present a rotamer change with respect to the apo structure when a TM-based compound is bound, while the site at Asp14–Met29 presents a change in the rotameric state of the methionine when the monodentate compound binds to it. In the case of the bidentate compound, both Asp14 and Met29 change the rotameric state upon binding. More details about the rotameric states, as well as the RMSD values, are reported in the SI (Table S2).

In the case of Metal3D, three regions with high probability are found consistently in all the structures: two of them in proximity of His119 either in the direction of His12 or Asp121, respectively, and one corresponding to a disulfide bridge between two cysteine residues (predictions for the apo structure in Fig. 6 and for all other structures in Fig. S4). Similar to what has been observed in the case of HEWLC, all four disulfide bridges in the structure are also identified by Metal3D, with a lower associated probability. The binding site at His105 is not predicted, with the exception of the PDB structure 4S18, in which this site is occupied by a Pt ion from oxaliplatin. The probability density at this site, however, has a low probability (p∼0.1) and is not consistent between the other structures, even if the side chain positioning of His105 varies only slightly among all structures without a change in the rotameric state. In particular, His105 presents an all-atom RMSD of ≤ 0.3 Å (a detailed comparison is given in Tab. S3 and Fig. S3). Furthermore, Metal3D does not predict one of the known binding sites involving Asp14 and Met29 (PDB 4S18 and 4S0Q). The reason for that might be the presence of a methionine at the site, which is an amino acid commonly bound by platinum, but not zinc (Fig. 2), on which Metal3D was trained. However, a low-probability region between Asp14 and His48 is identified in some of the structures, in close proximity to Met29.

**Figure 6:**
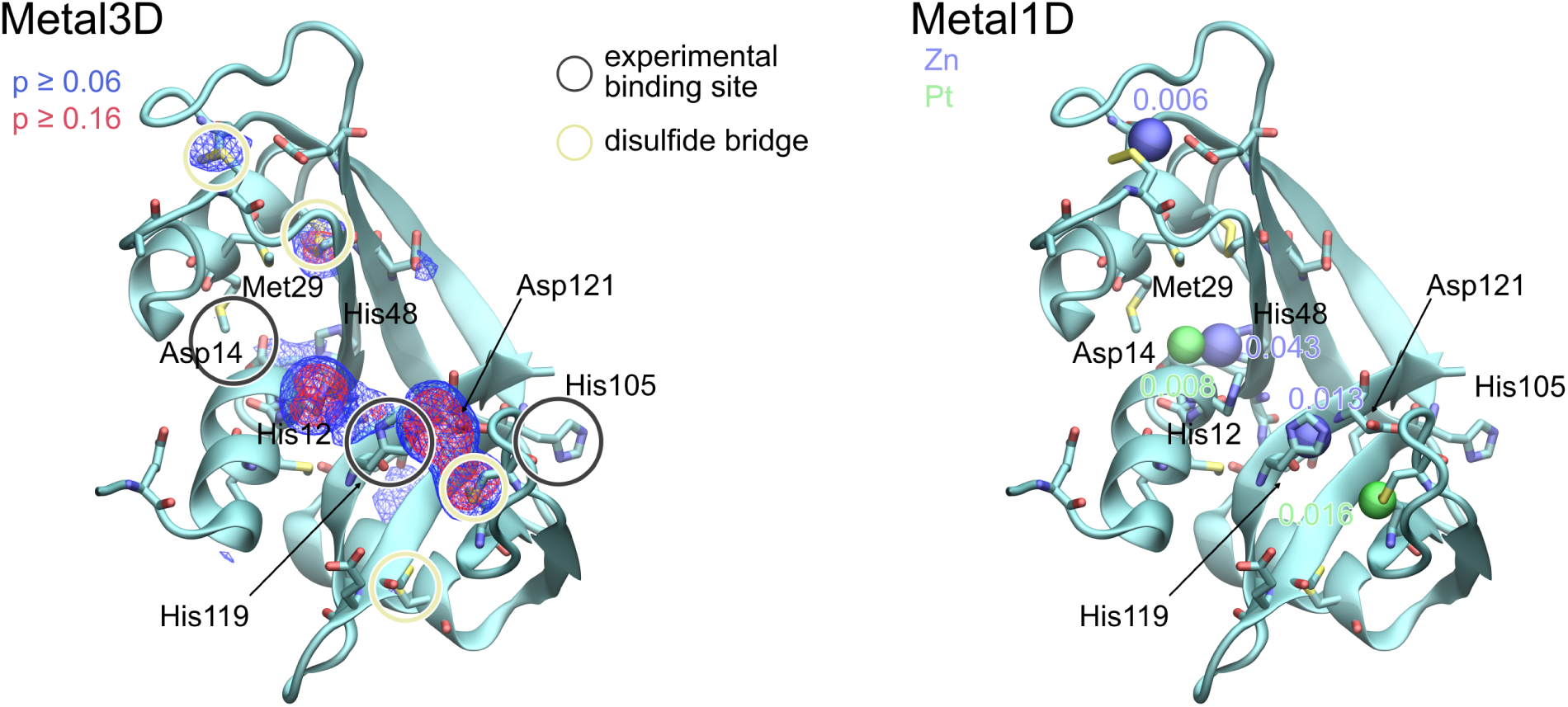
Metal3D (left) and Metal1D (right) predictions for RNaseA apo structure (PDB 1RPH). The experimental binding sites and disulfide bridges are highlighted with circles in the structure on the left. In the test set considered, Pt-based compounds bind all three sites, while the Ru-based compound binds at His105. For Metal3D, two probability density contours are represented in different colors as isosurfaces with a wireframe representation, in blue the one with lower probability and in red the one with higher probability (isovalues of 0.06 and 0.16, respectively). For Metal1D, the results obtained with different probability maps are reported: zinc (blue) and platinum (green), and for each prediction, the probability associated with each site is reported in the figure. For this structure, Metal1D did not predict any site for ruthenium binding.

In the case of Metal1D, the predictions based on the zinc probability map are in partial agreement with Metal3D: the His119 site is consistently identified in all the structures, except for one (PDB 5JLG), in which this residue assumes a different conformation (predictions for the apo structure in Fig. 6 and for all other structures in Fig. S5). Other consistent predictions among all of the structures correspond to the region with the disulfide bridge and to His48, in close proximity to Asp14, which together with Met29 is part of the bidentate binding site of oxaliplatin (PDB 4S18). However, also in this case, His105 is not identified as a possible binding site. In the case of the probability map generated from Pt-based compounds, the predictions partially agree with those based on zinc. Moreover, Metal1D predicts some disulfide bridges, but only in the case of the apo structure, and His48 in proximity to the Asp14–Met29 binding site is identified, similarly to what was observed for Metal3D. In the case of the Ru-based probability map, only some of the disulfide bridges are identified inconsistently between the different structures.

### 3.4 Dependence of the predictions on rotameric variations

#### 3.4.1 Monodentate histidine-containing sites

Based on the results for HEWLC and RNaseA discussed so far, it is possible to observe some dependence of the prediction on the rotamer conformations of the side chains of the metal binding amino acids. For both predictors, this is reasonable, considering that the local environment processed by the 3DCNN for Metal3D, and the geometric criteria for Metal1D, will be optimal when the conformation is close to that of a metal binding site. However, a large degree of dependency on the rotamer conformations is detrimental for the prediction of unknown binding sites starting from an apo structure, since it is possible to overlook some binding sites simply because the amino acids are in a different conformation.

To elucidate the dependency on rotamer state, we selected some of the binding sites in our test enzymes and probed the effect on the predictions upon variations of the rotamer state of one of the amino acids (Fig. 7). In particular, in the apo structures, we selected His15 in HEWLC and His105 in RNaseA as representative cases of a successful and unsuccessful prediction, respectively. For both structures, we investigated the influence of sampling different rotamer conformations on the prediction probabilities for Metal3D and Metal1D.

**Figure 7:**
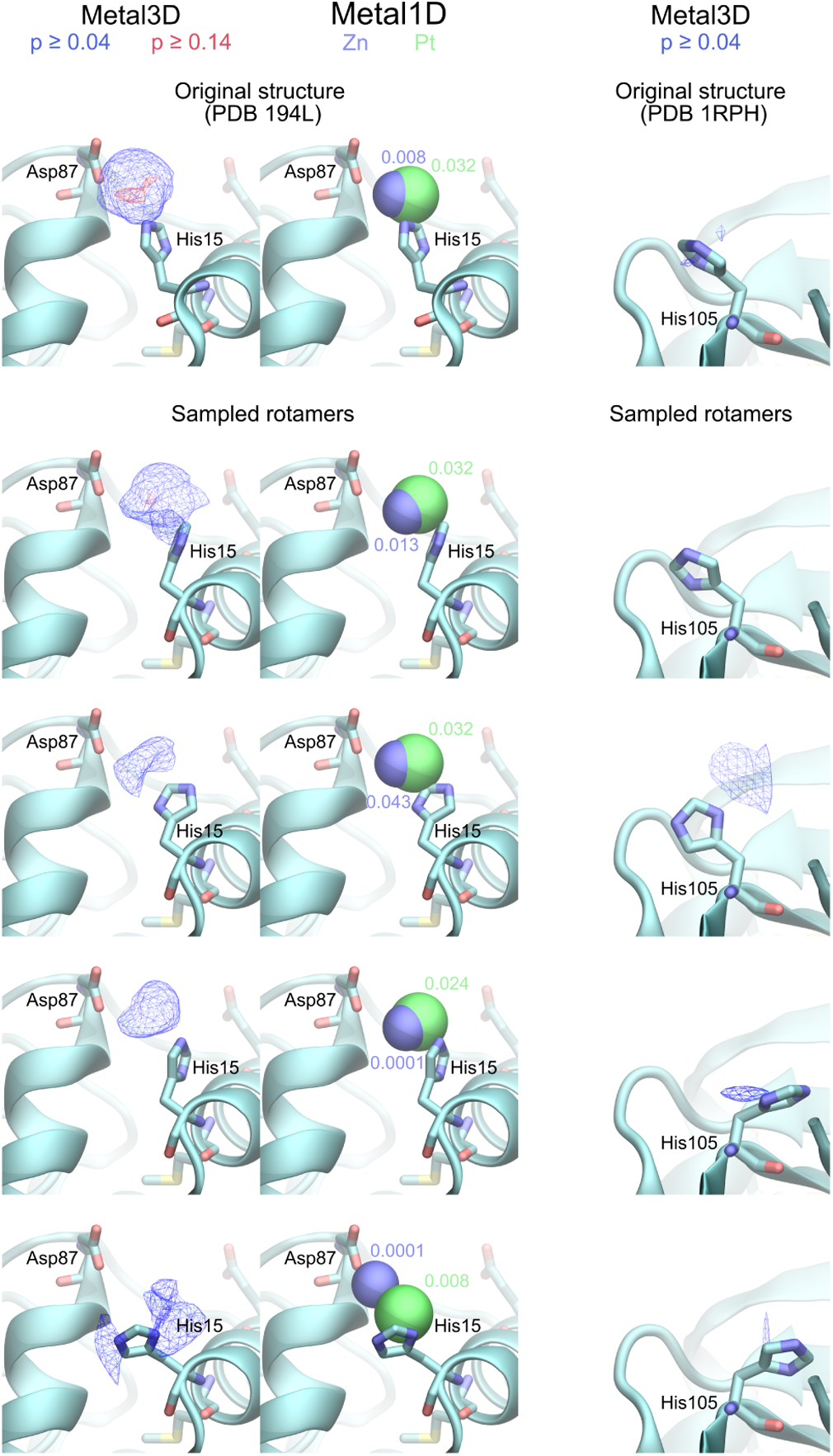
Metal3D and Metal1D predictions for different rotamers of His15 in HEWLC and His105 in RNaseA. In both cases, the initial structure is the apo structure, and only a single rotamer is changed. In the case of His105, Metal1D does not predict any site for the different rotamers, hence only Metal3D predictions are reported.

In the case of His15, the holo structure presents a different rotamer for the imidazole ring, but not a change in the orientation of the side chain. This site was already predicted with sufficient confidence in the apo structure, but the sampling of further rotamers results in a decrease of the Metal3D probability around the amino acid. In particular, the region with high probability identified in the apo structure (p∼0.14) disappears for all of the other rotamers, while the region with lower probability (p∼0.04) remains present, even if its shape and size change. In the case of the predictions with Metal1D, all the rotamers sampled lead to a prediction with the zinc and platinum probability maps in proximity to His15, in analogy to what was observed in the apo structure. The probability value is similar to that of the original structure, especially in the case of the predictions with the probability map for platinum. Similarly to what has been observed in the apo structure, no prediction is made with the probability map extracted from the Ru-based compounds. In the case of His105, which was not predicted for the apo structure, the sampling of different rotamers leads Metal3D to identify low-probability regions (p∼0.004) for some of the sampled rotamers. In contrast, Metal1D does not make any prediction in the proximity of His105 for any of the sampled configurations.

This comparison highlights some differences between the two predictors: on one hand, Metal3D seems to be more sensitive to the side chain positioning and the rotameric state, with significant differences in the probability regions generated when different rotamers are considered. However, none of the rotamers tested for His15 led to a significant increase in the probability with respect to the apo structure, in which a high probability region was already identified. On the other hand, in the case of Metal1D, the dependence on the rotameric state of the side chain seems to be less pronounced, and the probability values associated with the predicted metal site are more consistent in the case of the probability map extracted from Pt-based compounds. This observation is in line with what was observed in the original publication by Dürr et al.,^12^ where Metal3D presented similar precision and recall for TM ions different from zinc, while Metal1D showed similar precision, but lower recall.

#### 3.4.2 Bidentate glutamate-containing site

To further test the sensitivity to the rotamer conformations and the prediction ability of Metal1D and Metal3D, we also performed additional predictions for a known bidentate TM binding site in NCPs containing two glutamates (GLU site). For our work, this represents an interesting system considering that the two glutamates significantly change their rotameric state when a metal compound binds. Fig. 8 shows the GLU site from the apo and holo structures, from PDB ID 3REH^34^ and 5XF3,^36^ respectively, as well as the predictions from Metal3D and Metal1D. Only in the case of the holo structure, in which the glutamates are in the conformation corresponding to the binding of the metal compound, the predictors identify the site. In the case of Metal3D, a high probability region with p∼0.2 can clearly be identified, i.e., with a significantly larger probability than for the sites tested so far, possibly thanks to the bidentate nature of the site. For Metal1D, the predictions with the three probability maps (zinc, platinum, and ruthenium) are all able to locate this binding site. However, in the apo structure, no prediction is made by either Metal3D or Metal1D.

**Figure 8:**
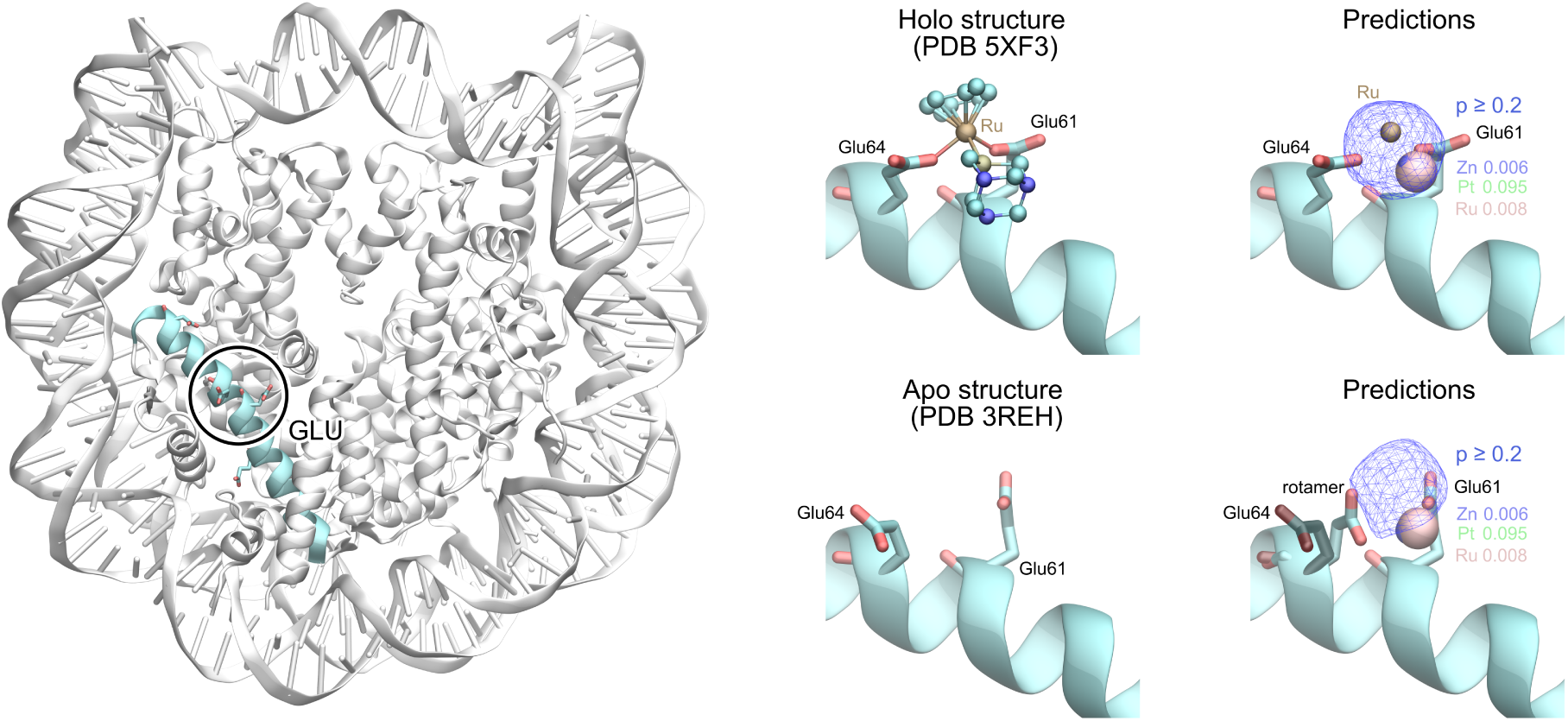
Metal3D and Metal1D predictions for the GLU site in the case of the holo (top) and apo (bottom) structures. For the latter, we also report the successful prediction after sampling a different rotamer for Glu64. In the case of Metal1D predictions, the metal ions located using the three different probability maps overlap, and only the one corresponding to ruthenium is visible, but the corresponding probability values for each probability map are indicated.

When testing the influence of different rotamer conformations for the two glutamates in the GLU site, in the case of Glu61, none of the tested rotamers leads to a favorable conformation for metal binding (Fig. S6). As a result, neither Metal3D nor Metal1D identifies a binding region between Glu61 and Glu64. However, in the case of Glu64, some of the rotamers assume a ‘favorable’ conformation, pointing in the direction of Glu61. In particular, we report in Fig. 8 the corresponding predictions for one of such rotamers, which lead to similar predictions as the holo structure with the bound drug. Similarly to what is observed for the holo structure, Metal3D identifies a high probability region (p∼0.2), and Metal1D locates a metal ion with all three probability maps tested.

This last comparison highlights the influence of rotamer sampling for bidentate sites, in which it is reasonable to assume that sampling different rotameric states from the apo structure is more likely to lead to a conformation in which two (or more) amino acids are in a favorable conformation for metal site binding, which is not necessarily favorable in the absence of a TM ion.

## 4 Conclusion

In this work, we explored the applicability of the Metal3D and Metal1D predictors^12^ to identify the binding sites of TM-based compounds. Although they were originally designed to predict binding sites of zinc ions, our results demonstrate a surprising degree of predictive power for more complex TM binding sites. Both methods consistently identify some of the drug binding sites across the tested structures, enabling the identification of the most common binding sites from the apo conformations. However, their performance is limited in cases where the binding site involves only a single solvent-exposed residue.

Both predictors, particularly Metal3D, show sensitivity to the side chain positioning and rotameric state of the amino acids at the binding site. This dependence suggests that binding sites may be overlooked in apo structures if the key amino acids are not ‘pre-organized’ in a favorable orientation. To investigate this, we sampled alternative rotameric states for selected residues. For monodentate sites, where the TM-based compound binds a single amino acid, these variations had little impact on prediction outcomes, regardless of whether the original prediction was correct or incorrect. In contrast, for a bidentate site, introducing alternative rotamers enabled the correct identification of the binding site directly from the apo structure.

Overall, our findings suggest that Metal3D and Metal1D can serve as a first step in establishing rapid, low-cost pre-screening tools to highlight putative ‘hotspot’ regions for the binding of TM-based compounds. These predictions can then be refined by more accurate methods, such as QM/MM simulations. To design a robust pipeline for predicting unknown binding sites in apo structures, however, several limitations must be addressed. In particular, a clear difference between a TM ion and a TM-based compound is the additional steric volume occupied by the additional ligands coordinating the metal center in the TM-based compound. This could be taken into account, e.g., in Metal3D by extending the input channels of the 3DCNN to consider unoccupied voxels or, in the case of Metal1D, by adapting the geometric criteria. Moreover, in this study, we highlighted the dependence on the side chain orientation, which is a challenge partially inherent to any method based on single static structures. Possible improvements may come from integrating the predictors with some form of conformational sampling, e.g., through Monte Carlo methods or genetic algorithms optimization. Both approaches pose non-trivial challenges, as in Monte Carlo, suitable trial moves need to be established, balancing the increase in the predictor probability with maintaining energetically favorable conformations. In the case of genetic algorithms, a possible solution includes the use of multi-objective optimization^43^ by designing a fitness function based on the probability from the predictors and combining it with one for the stability of the resulting conformation. This second fitness function should describe the protein stability, taking into account the protein–metal interaction, but this cannot be easily evaluated, e.g., with classical FFs, due to their limitations in describing TMs.

Another crucial limitation is the dataset availability. For Metal1D, we generated probability maps from Pt- and Ru-containing structures, but their number is extremely small compared, e.g., to Zn. We observed some advantages in the case of the Pt-based probability map, but the Ru-based one was unsuccessful, possibly due to the extremely limited set of structures available to generate it. Expanding the dataset, potentially combining related TMs together, could improve the prediction accuracy and even provide sufficient data to train ML models, such as Metal3D.

In conclusion, while not yet fully general, Metal3D and Metal1D represent a valuable first step toward efficient prediction of TM compound binding sites. Further work will focus on addressing the limitations identified, with the ultimate goal of developing robust predictors capable of identifying unknown binding sites for TM-based compounds in proteins.

## Author Information

### Author Contributions

**AL**: conceptualization (equal), software (lead), investigation (lead), visualization (lead), writing — original draft (lead), writing — review & editing (equal). **UR**: conceptualization (equal), writing — review & editing (equal), supervision (lead), funding acquisition (lead).

### Notes

The authors declare no competing financial interest.

## Acknowledgments

This work has been supported by the Swiss National Science Foundation (grant 200020-185092 and 200020-219440).

## Data Availability

All the predictions for the different protein structures are deposited on Zenodo under https://zenodo.org/records/18416988, including VMD visualization states to reproduce the figures presented in the paper. In the same repository, we also included a Jupyter Notebook to perform all predictions with Metal3D and Metal1D, as well as generate the new probability maps for Metal1D.

## Supporting Information Available

Supporting Information contains detailed comparisons of the crystal structures used in the different test sets, as well as predictions with Metal3D and Metal1D for the holo structures not shown in the main text.

## Supporting Information

### Lysozyme C TM binding site prediction

**Table S1:** RMSD values for the different X-ray structures considered for the HEWLC. All structures have been aligned to the backbone of the apo structure (PDB ID 194L). The RMSD corresponding to an occupied binding site is indicated in bold, and an asterisk indicates a rotamer change with respect to the reference (apo) structure.

| PDB ID | RMSD (Å) |  |  |  |  |
| --- | --- | --- | --- | --- | --- |
|  | backbone | metal binding<br>amino acids | binding sites |  |  |
|  |  |  | His15 | Asp101 | Asp119 |
| 194L | ref. | ref. | ref. | ref. | ref. |
| 2I6Z | 0.25 | 0.38 | **1.14\*** | 0.42 | 0.22 |
| 5II3 | 0.25 | 0.53 | **1.18\*** | 1.62* | 0.37 |
| 5V4G | 0.28 | 0.54 | **1.19\*** | **1.68\*** | 0.12 |
| 6QEA | 0.21 | 0.54 | **1.15\*** | 1.69* | 0.25 |
| 6WGO | 0.31 | 0.91 | 0.24 | 3.83* | **0.45** |

**Figure S1:**
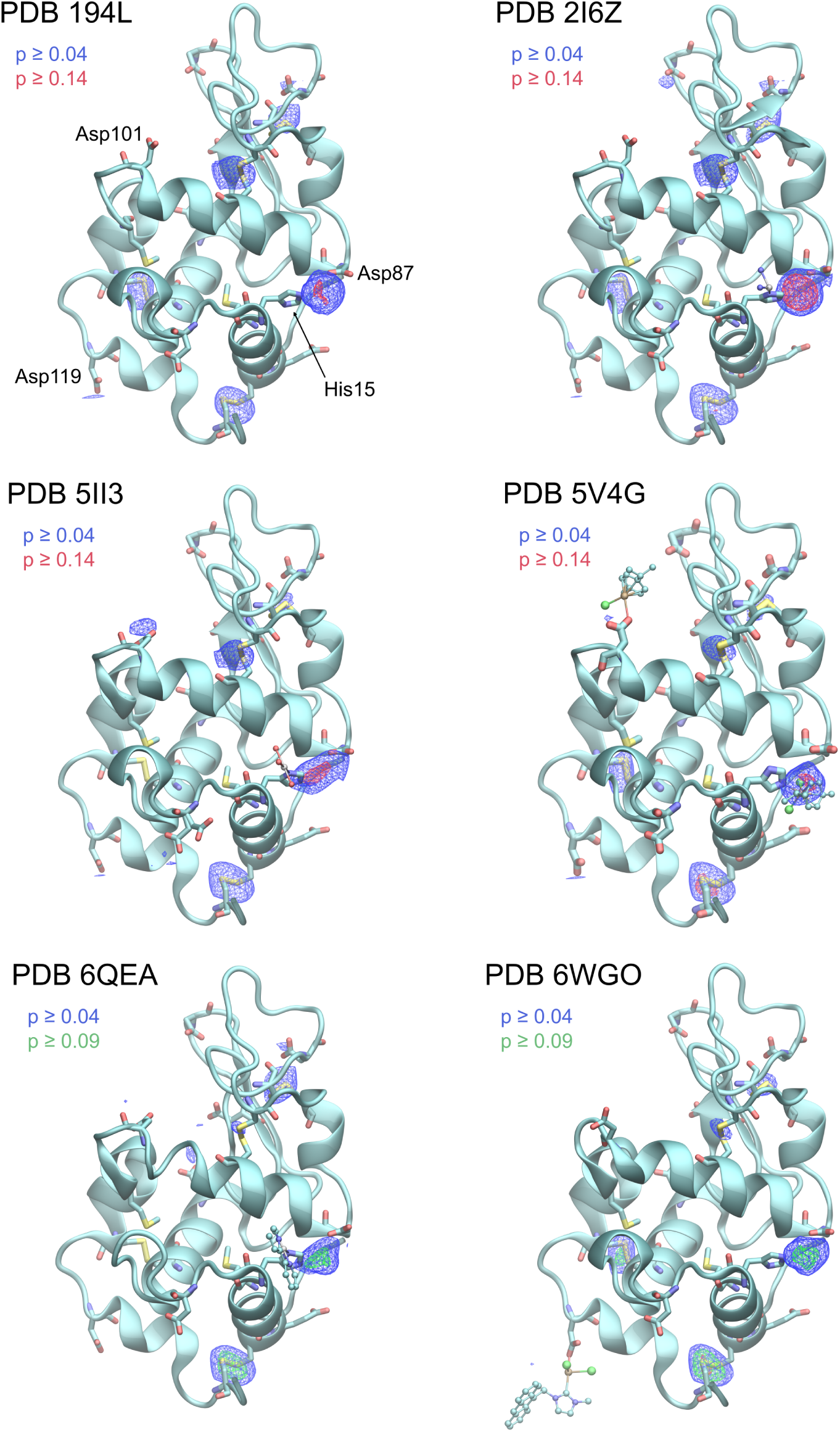
Metal3D predictions for HEWLC made on the different crystal structures. Metal3D probability densities are represented in different colors as isosurfaces with a wireframe representation. The isovalue for the low probability predictions (blue) is 0.04, while the one for higher probability predictions (red) is 0.14. In the cases where no region had probability higher than 0.14, a lower isovalue is used (0.09, green).

**Figure S2:**
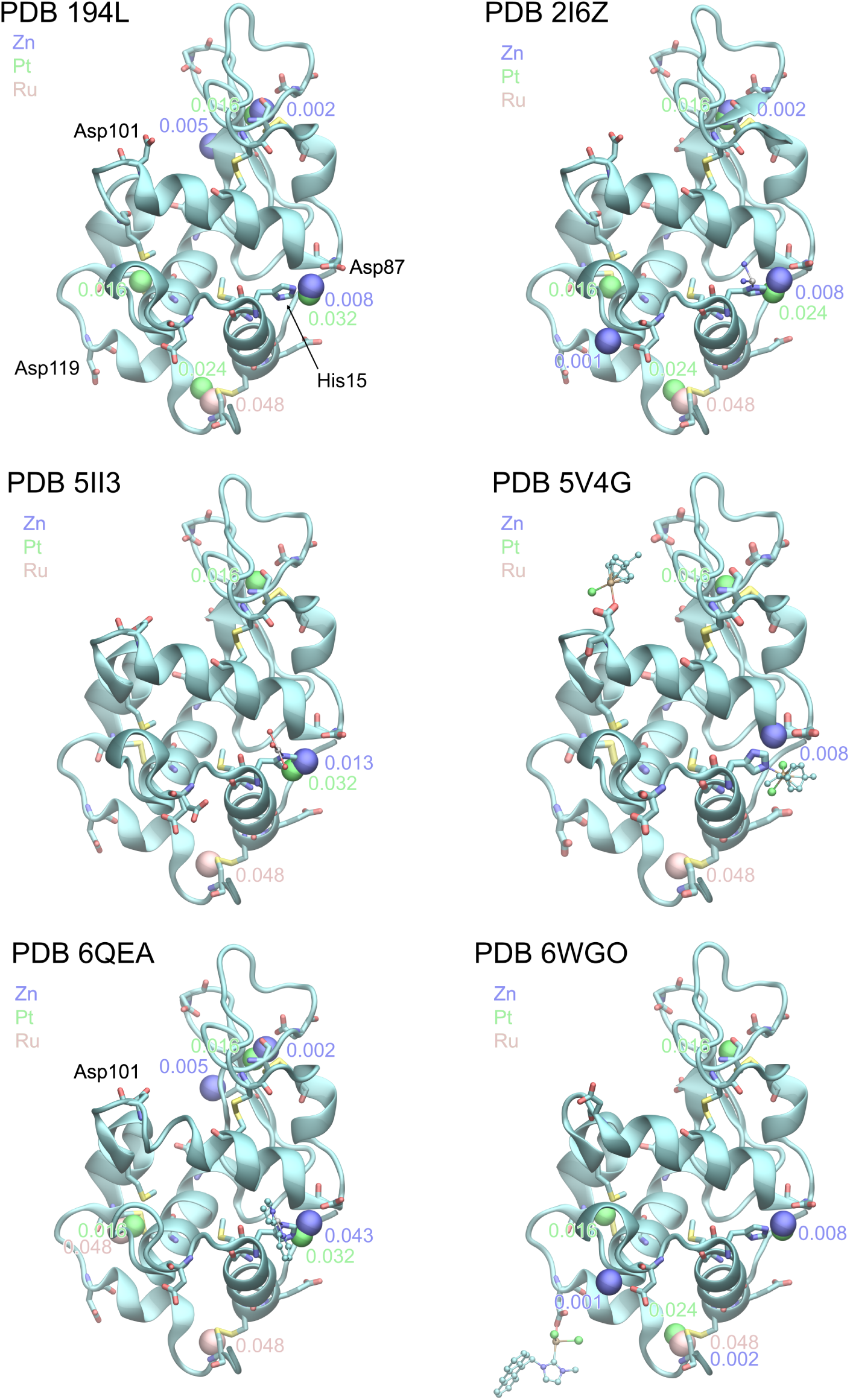
Metal1D predictions for HEWLC made on the different crystal structures with different probability maps: zinc (blue), platinum (green), and ruthenium (pink). For each prediction, the probability associated with each site is reported with the same color as the corresponding probability map.

### Ribonuclease A TM binding site predictions

**Table S2:** RMSD values for the different X-ray structures considered for the RNaseA enzyme. All structures have been aligned to the backbone of the apo structure (PDB ID 19PH). The RMSD corresponding to an occupied binding site is indicated in bold, and an asterisk indicates a rotamer change with respect to the reference (apo) structure.

| PDB ID | backbone | metal binding<br>amino acids | RMSD (Å) |  |  |
| --- | --- | --- | --- | --- | --- |
|  |  |  | binding sites |  |  |
|  |  |  | Asp14/Met29 | His105 | His119 |
| 1RPH | ref. | ref. | ref. | ref. | ref. |
| 4S0Q | 0.51 | 0.68 | 0.79/ **0.91** * | 0.38 | 0.44 |
| 4S18 | 0.48 | 0.54 | **0.72** */ **1.1** * | **0.29** | **0.33** |
| 5JLG | 0.52 | 0.48 | 0.18/0.40 | **0.38** | 3.09* |

**Table S3:** RMSD values for the His105 residue in the different X-ray structures considered for the RNaseA enzyme. All structures have been aligned to the backbone of the apo structure (PDB ID 1RPH) and the RMSD is computed for all the atoms of His105 using the structure of PDB 4S18 as reference, for which Metal3D made a successful prediction.

| PDB ID | RMSD (Å) – His105 |
| --- | --- |
| 1RPH | 0.29 |
| 4S0Q | 0.15 |
| 4S18 | ref. |
| 5JLG | 0.16 |

**Figure S3:**
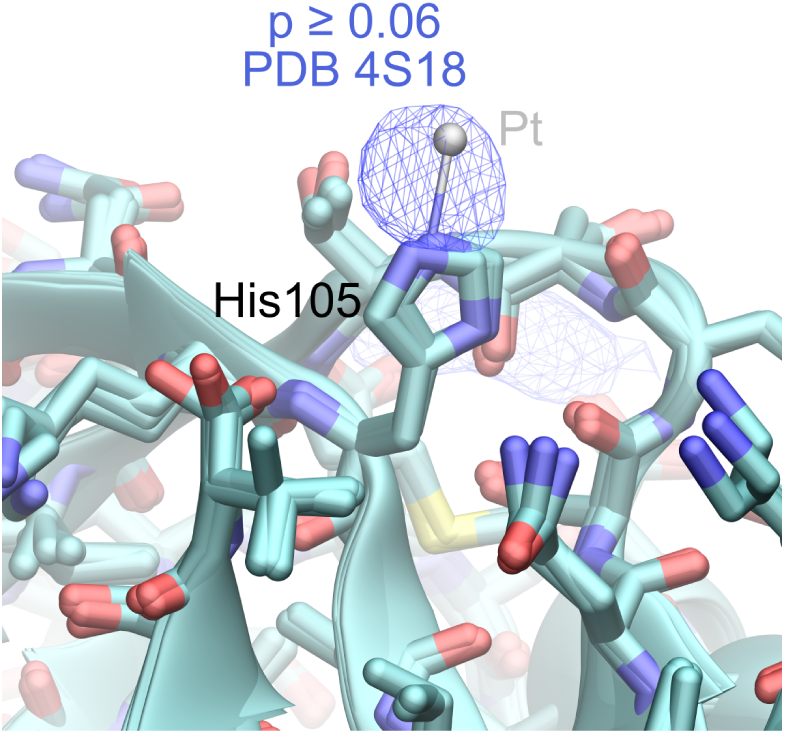
Comparison of the environment around the His105 residue. All structures have been aligned to the backbone of the apo structure (PDB ID 19PH), and the probability density generated from Metal3D for PDB 4S18 is also represented in blue.

**Figure S4:**
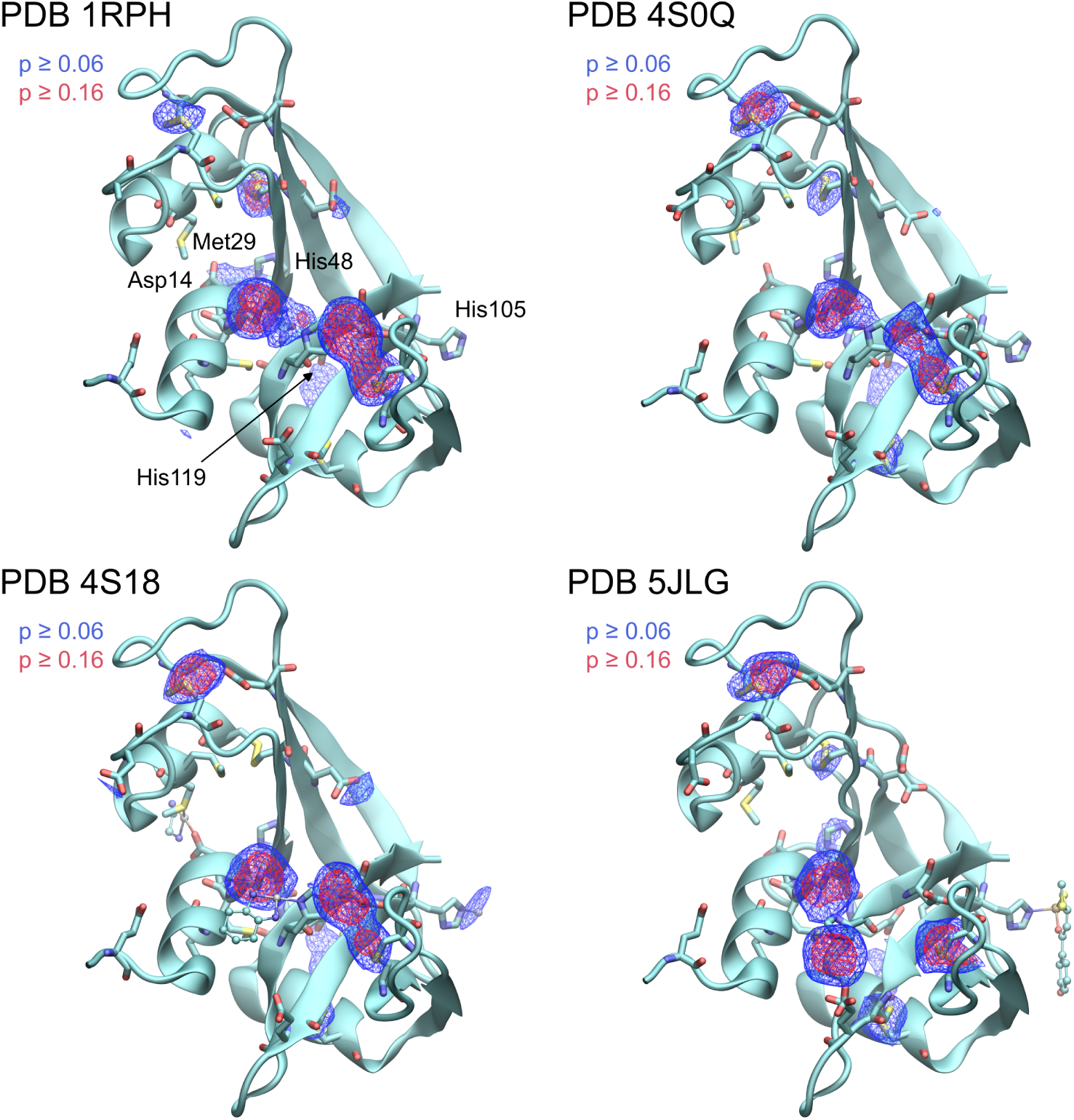
Metal3D predictions for RNaseA made on the different crystal structures. Metal3D probability densities are represented in different colors as isosurfaces with a wireframe representation. The isovalue for the low probability predictions (blue) is 0.06, while the one for higher probability predictions (red) is 0.16.

**Figure S5:**
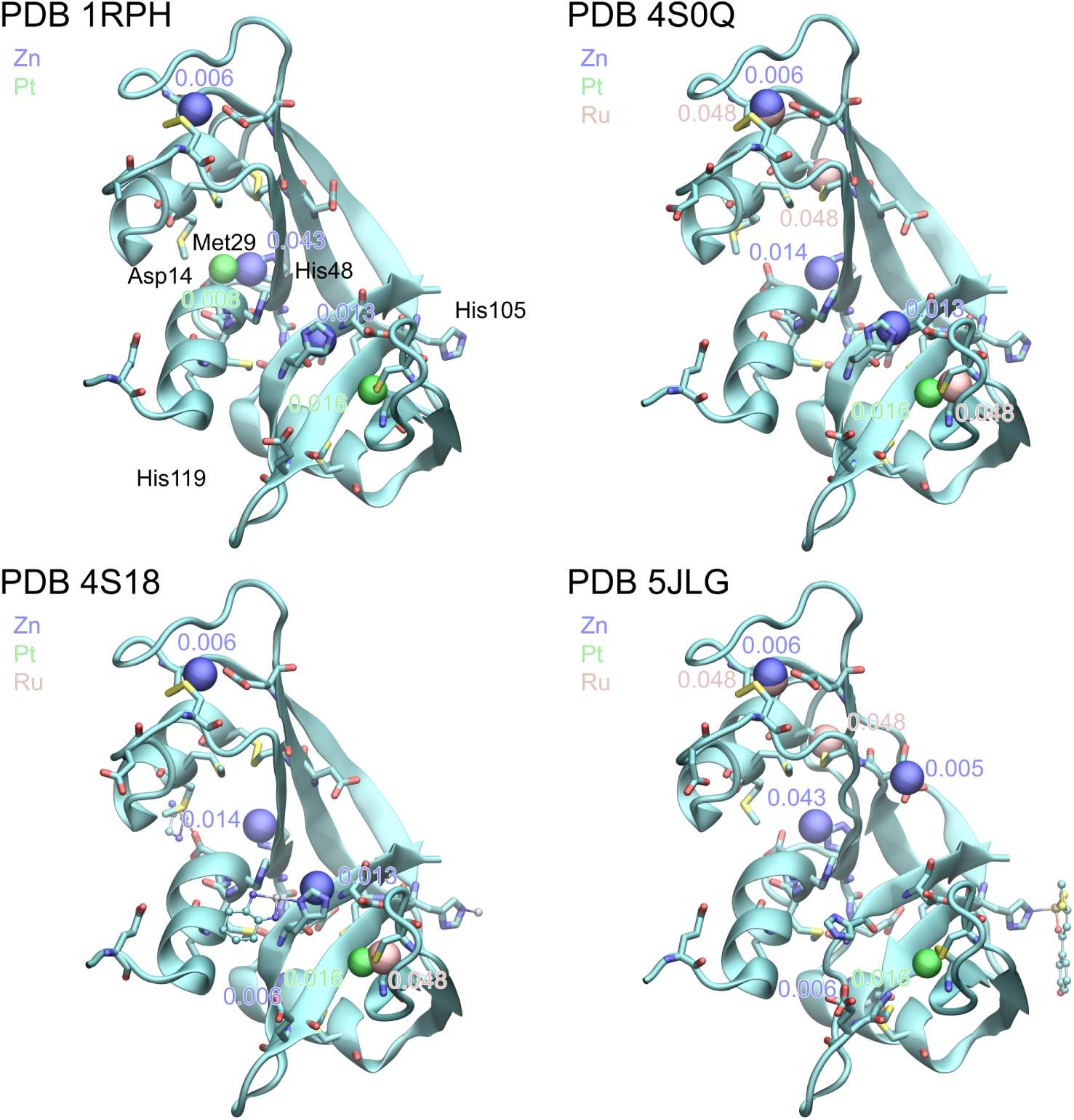
Metal1D predictions for RNaseA made on the different crystal structures with different probability maps: zinc (blue), platinum (green), and ruthenium (pink). For each prediction, the probability associated with each site is reported with the same color as the corresponding probability map.

### Dependence of the predictions on rotamer conformation

**Figure S6:**
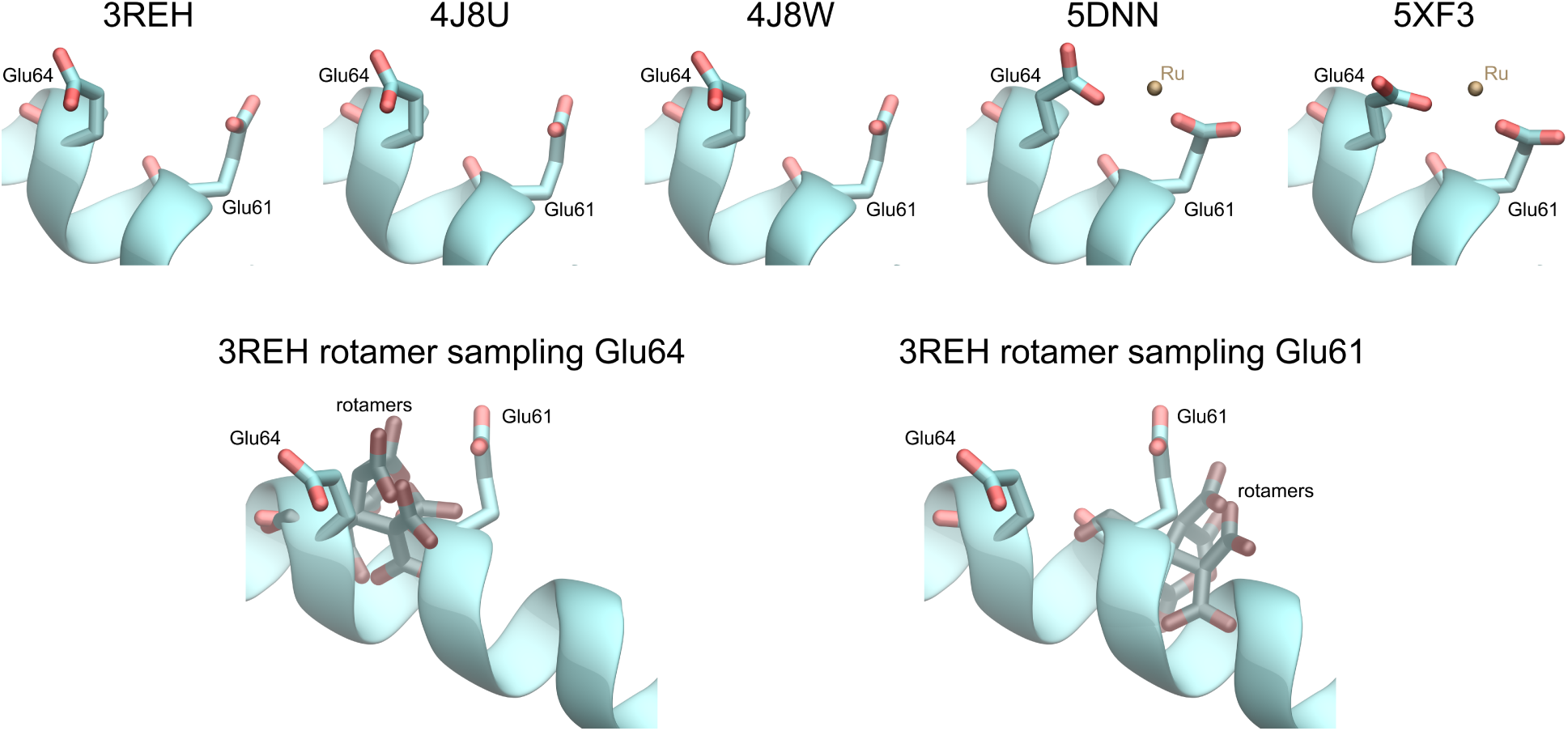
GLU site conformations for the apo and holo protein from different PDB structures. For the apo structure (PDB ID 3REH), the different sampled rotamers for Glu61 and Glu61 are also reported.

